# Treatment of Graft-versus-Host Disease by Echinomycin in a New Humanized Mouse Model

**DOI:** 10.1101/108951

**Authors:** Yan Liu, Christopher Bailey, Christopher Lazarski, Chun-shu Wong, Pan Zheng, Yang Liu, Yin Wang

**Affiliations:** Center for Cancer and Immunology Research, Children’s Research Institute, Children’s National Medical Center, Department of Pediatrics George, Washington University School of Medicine Washington, DC 20010, USA

## Abstract

Drug development effort against GVHD is hampered by the lack of clinically relevant humanized animal models for preclinical testing. Current humanized GVHD models rely on adoptive transfer of a high number of human peripheral blood mononuclear cells (PBMCs) into immunodeficient mice. Here we report a novel humanized GVHD model by transplanting a small number of human BM cells into newborn *NOD. SCID IL2ry*^0^ (NSG) mice. Transplantation of human BM cells (BMT) causes acute GVHD, with lethality between 15 to 60 days. Pervasive human T-cell infiltration into multiple organs, including lung, intestine, skin, kidney, liver, and stomach, was observed in all mice analyzed. Surprisingly, the human T cells express high levels of hypoxia inducible factor 1α (HIF1α) protein even under normoxic environment. Administration of Echinomycin, a potent inhibitor for HIF1α, rapidly ablated HIF1α protein in T cells and gradually reduced the frequency of human cells in the peripheral blood and target organs. Echinomycin provides a sustained therapeutic effect, as demonstrated by dramatic reduction of clinical symptoms, pathology score and by doubling of the median life span of the chimeric mice. Our results reveal a critical role of HIF1α in GVHD and demonstrate that HIF1α inhibitors such as Echinomycin should be explored for clinical drug development against GVHD.

## Introduction

Allogeneic hematopoietic stem cell transplantation (HSCT) is a potential curative therapy for hematologic malignancies. While the lymphocytes in donor bone marrow (BM) play a critical role in the prevention of tumor relapse, they are also responsible for the development of graft-versus-host disease (GVHD). GVHD causes multi-organ damage and is one of the leading causes of morbidity and mortality associated with HSCT in patients^1–3^. Despite advances in preventing GVHD in humans by the use of non-specific immunosuppressive drugs, GVHD remains a significant cause of morbidity and mortality following HSCT ^4–6^.

A clinically relevant humanized animal model is critical for the development of new strategies for the treatment of GVHD. Significant advances have been made towards generating humanized GVHD in mice^7–11^. However, these models have significant drawbacks. To date, the best models take advantage of immunodeficient mice, particularly the Non-obese diabetic (NOD). *Scid. IL2r*γ^*0*^ (NSG) strain, in which transplantation of human PBMC is performed via intravenous route in adult mice^7,8^. Although these models exhibit lethal GVHD, they do not fully recapitulate pathogenesis of the human disease. For example, the immune damage is notably most severe in the lung, while only mild infiltration to the skin and liver is achieved ^7,8^, the latter being the most common target organs in human GVHD ^2^. Furthermore, all current models that rely either on human PBMC or T-cell purified cord blood ^12^ to induce xeno-GVHD do not address the fact that human GVHD occurs following HSCT in which BM, rather than peripheral blood, is the main source of HSC. It is unclear if T cells in the BM and peripheral blood respond similarly to therapies.

During the effector phase of the GVHD response, donor T cells recognize antigens in the recipient as foreign, resulting in the expansion of donor T cells and immune engagement upon host tissues. Identification of druggable molecular targets may help to treat GVHD. In this context, it has been demonstrated that hypoxia-inducible factor 1α (HIF1α) plays a critical role in driving T cell differentiation, metabolism and cytotoxic activity ^13–15^. Furthermore, it has been shown that activation of T cells both induces and stabilizes HIF1α, and that the stabilized HIF1α was responsible for increasing the cytolytic activity of CD8^+^ by expression and release of cytolytic molecules and costimulatory molecules ^14–16^. It has been reported that HIF1α binds to Foxp3 and induces its degradation, and thereby inhibits regulatory T cell development ^17,18^. Deletion of *Hif1a* restores Foxp3 expression and rescues the defective suppressive activity in *Dtx1*^*-/-*^ regulatory T cells ^19^. Given the multiple functions of HIF1α that can potentially impact GVHD pathogenesis, we set out to evaluate its contribution to GVHD.

Here, we report the establishment of a novel humanized GVHD model by transplanting a minute amount (0.5 × 10^6^ cells) of human bone marrow cells (BMT) into newborn NSG mice. The BMT reproducibly induces a systemic acute GVHD syndrome that faithfully recapitulates pathological features of human GVHD. Surprisingly, the human T cells accumulated HIF1α under normoxic environment much alike what we have reported in leukemia stem cells ^20^. Treatment with Echinomycin, a potent HIF inhibitor, provides a sustained therapeutic response. Our data demonstrate HIF1α as a novel therapeutic target for GVHD.

## Materials and Methods

### Mice

All procedures involving experimental animals were approved by Institutional Animal Care and Use Committees of the Children’s Research Institute where this work was performed. *Nod. Scid. Il2rg^0^* (NSG) mice were purchased from the Jackson Laboratory and were bred and maintained under specified pathogen-free conditions in research animal facilities at the Children’s Research Institute.

### Xenografting of human BM cells into newborn NSG pups

Human bone marrow mononuclear cells isolated from healthy human adult bone marrow using density gradient separation were purchased from Stemcell Technologies (Vancouver, Canada) and Lonza (Walkersville, MD, USA). Human CD3^+^ cells from BM were sorted from human BM cells using BD Influx (BD Biosciences). Cells were thawed, counted and re-suspended in 1XPBS in a concentration of 0.1-0.5×10^6^ per 30 µl. 0.1-0.5×10^6^ cells were transplanted via intrahepatic injection into irradiated (1.30 Gy) newborn NSG pups. Human CD45^+^ cells in PBMC of recipients were detected by FACS analysis at day 17 to 20 after transplantation.

### Flow Cytometry

Peripheral blood was collected by sub-mandibular bleeding at different times after transplantation of human BM cells. Fluorochrome-labeled antibodies were directly added into whole blood. After 30 min of staining, the unbound antibodies were washed away and the red blood cells were lysed with BD FACS™. The stained cells were analyzed with BD FACS Canto II flow cytometry.

Spleens were gently grinded with frosted objective slides and bone marrow was dissociated with syringes to obtain single-cell suspensions and passed through a nylon cell strainer, washed three times with RPMI-1640, labeled with antibodies and analyzed for the presence of different human cell populations.

Antibodies used were phycoerythrin (PE) conjugated anti-human CD45, anti-human Foxp3, PE-Cy7 conjugated anti-human CD4, allophycocyanin (APC) conjugated anti-human CD11c, anti-human Foxp3, anti-mouse CD45, eFluor 450 conjugated anti– human CD3, eFluor 780 conjugated anti-human CD19 (eBioscience, San Diego, CA), peridinin chlorophyll protein conjugated (PerCP) anti–human CD14, fluoresceinisothiocyanate (FITC) conjugated anti-human CD8, V500 conjugated anti-human CD14 (BD Bioscience, San Jose, CA), and PE-Cy 7 conjugated anti-human CD11b (BioLegend). We also used PE, PerCP and APC conjugated anti-human HIF1α from RD Systems (Minneapolis, NM).

### Immunohistochemistry

Immunohistochemistry was performed on tissue sections from skin, liver, lung, spleen, and kidney of humanized NSG mice. Sections were fixed with 4% paraformaldehyde and dehydrated with graded alcohol. After treatment with heated citrate buffer for antigen retrieval, sections were blocked for endogenous peroxidase activity. Sections were then incubated in 10% goat serum followed by primary antibodies at 4°C overnight. Fixed samples were stained with the following antibodies, anti–human CD45 (MEM-28) and CD3 (ab828, abcam), for detection of infiltrated human cells. After incubation for 30 minutes with secondary antibody, the specimens were visualized by DAB treatment. Sections were lightly counterstained with hematoxylin to enable visualization of nuclei.

### Immunofluorescence

After deparaffinization and rehydration of slides with xylene and ethanol, tissue sections from mice were treated with 10 mM sodium citrate buffer, pH 6.0. The sections were permeabilized with 0.3% Triton X-100 in 10 mM Tris-HCl buffer for 30 min. After blocking with 2% bovine serum albumin (BSA) for 60 min, sections were incubated with primary antibody diluted in 10 mM Tris-HCl buffer containing 2% BSA at 4ºC, overnight, with subsequent staining with secondary antibody in BSA-Tris-HCl buffer at room temperature for 2-4 hrs. The nuclei were stained with DAPI. Slides were mounted with Prolong Antifade mounting buffer (Invitrogen, Carlsbad, CA 92008). Antibodies for hCD3 (NBP1, Novus), hCD45 (MEM-28, abcam), hCD4 (BC/1F6, abcam), and hCD8 (SP16, abcam) were used for immunofluorescence.

### Pathology scores

The organs of moribund mice were fixed in formalin and sectioned for hemoxylin and eosin (H&E) staining. All sections were scored based on the following criteria. Grade 0, no lesions; grade 1, minimal perivascular leukocyte infiltrations; Grade 2, Mild perivascular leukocyte infiltations; Grade 3, moderate perivascular leukocyte infiltrations, with leukocyte infiltration into parenchyma, tissue cell necrosis; Grade 4, Moderate to severe perivascular leukocyte infiltration, with intra-parenchymal leukocytes and tissue cell necrosis.

### Treatment of GVHD mice with Echinomycin

Newborn NSG pups received 0.3-0.5×10^6^ human BM cells intrahepatically. Beginning on day 17 or 27, the recipients received 5 intraperitoneal injections of Echinomycin, daily at 10 μg/kg. Following by 2 days of rest, and completed with another 5 intraperitoneal injections of Echinomycin at the same dose, once per day. The second round of treatment was performed following 5 days of rest from the prior round of treatment, using the same dose once every other day for 10 total treatments.

## Results

### Transplantation of human BM cells into newborn NSG pups causes GVHD

To develop a robust mouse model for human GVHD, we transplanted 0.5 × 10^6^ human BM cells into newborn NSG mice and followed the engraftment and expansion of human leukocytes, as well as survival of the chimera mice. The data for donor 1 are presented in Fig 1 and those for donors 2-4 are presented in Fig. S1.

**Fig.1.**
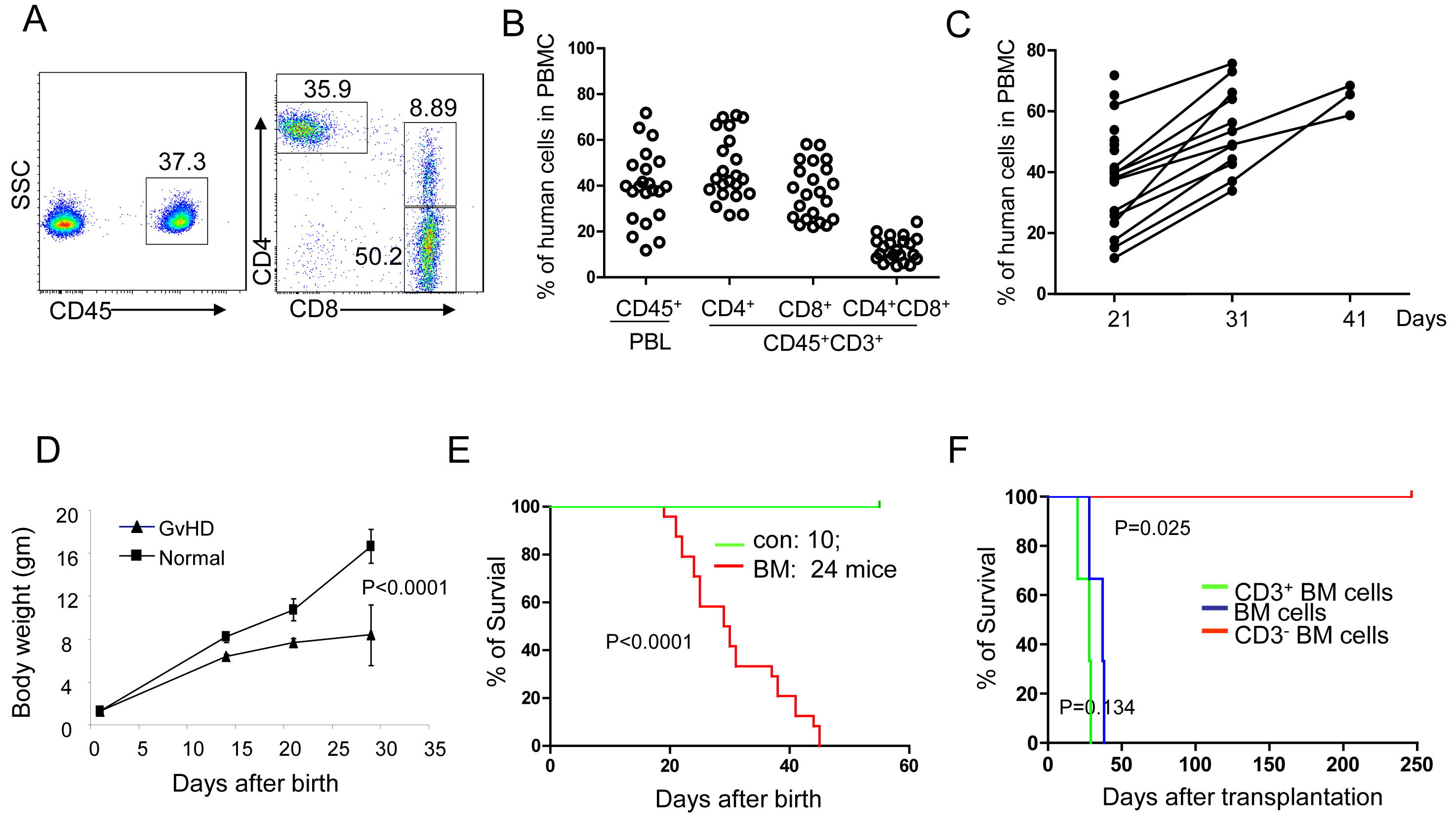
Transplantation of human BM cells into newborn NSG mice causes acute GVHD. Newborn NSG pups irradiated with 1.3 Gys were given intrahepatic injection of 0.5X10^6^ human BM cells. A. Representative FACS profiles depicting distribution of human CD45^+^, CD4^+^, CD8^+^, and CD4^+^CD8^+^ in PBMC of recipient at day 17 post-transplantation. B. Summary of percentage of human CD45^+^, CD45^+^CD4^+^, CD45^+^CD8^+^, and CD45^+^CD4^+^CD8^+^ in PBMC of recipients at day 17 post-transplantation. C. Longitudinal analysis for expansion of human cells in the NSG mice. Data shown are percentage of human CD45^+^ cells in PBL of each recipient at days 21, 31, and 41 were measured by FACS analysis. D. The bodyweight of irradiated (1.3Gy) normal NSG mice (n=25) and those that received irradiation and intrahepatic injection of 0.5X10^6^ human BM cells (GVHD, n=30) at birth. Data shown were means and SEM. E. Kaplan-Meier survival curves of human BM NSG recipients transplanted with BM cells (n=24) and normal NSG mice (n=10). Mice were observed daily for survival. Control mice (green curve) received irradiation only. F. Kaplan-Meier survival curves of human BM NSG recipients transplanted with human BM cells, sorted CD3^+^ and CD3^-^ cells from human BM cells. Mice were observed daily for survival.

As illustrated Fig. 1A and summarized in Fig. 1B, robust engraftment and expansion of human leukocytes were observed in the blood of the recipient mice on day 17, as an average of 40% human leukocytes were observed in the PBL. Among them, the overwhelming majority expressed T cell markers CD4 and/or CD8. As reported by others, approximately 10% of human T cells express both CD4 and CD8, although the function of this subset is unclear. Longitudinal studies showed gradual expansion of human cells until they surpassed mouse leukocytes (Fig. 1C). The first sign of GVHD is observable within two weeks, as judged by retarded growth (Fig. 1D) and damage to the skin and lack of hair growth (data not shown). Morbidity and/or mortality were observed starting on the third week, and essentially all mice die within six weeks (Fig. 1E). Similar engraftment and mortality were observed with other donors tested (Supplemental Fig. S1).

To test if human CD3^+^ caused GVHD in the recipient mice, we sorted human BM cells into CD3^+^ and CD3^−^ populations and transplanted the same number of either CD3^+^, CD3^-^, or unsorted bone marrow cells (BMC). As shown in Fig. 1F, recipients of CD3^+^ cells, as well as unsorted BMC, developed severe GVHD and died within 30 days. Since the GVHD onset was faster in the recipients, T cells were likely responsible for GVHD. In support of this notion, recipients of CD3^-^ cells never developed GVHD during our observation time of 246 days (Fig. 1F). Thus, transplantation of a very low number of human BM cells into newborn NSG pups causes severe xenogeneic GVHD, and CD3^+^ cells contribute to the GVHD in our humanized mouse model.

### Pathological features of the xenograft BMT-GVHD

Based on H&E staining, the BMT recipients have extensive inflammation in multiple organs (Fig. 2A, and Table 1). We performed immunohistochemical examination of the spleen, skin, liver, kidney, lung, pancreas, stomach and intestine of mice developing clinical symptoms of xenogeneic GVHD and detected human CD3^+^ T cells in these tissues (Fig.2B). Infiltration by human T cells was highest in the spleen and lungs, although high degree of infiltration was also apparent in the other tissues. In the lung, there were multifocal aggregates of human cells that expanded into the alveolar septa. In addition, we observed dramatic thickening of the skin, accompanied by human T cell infiltration along the dermal-epidermal junction.

**Fig.2.**
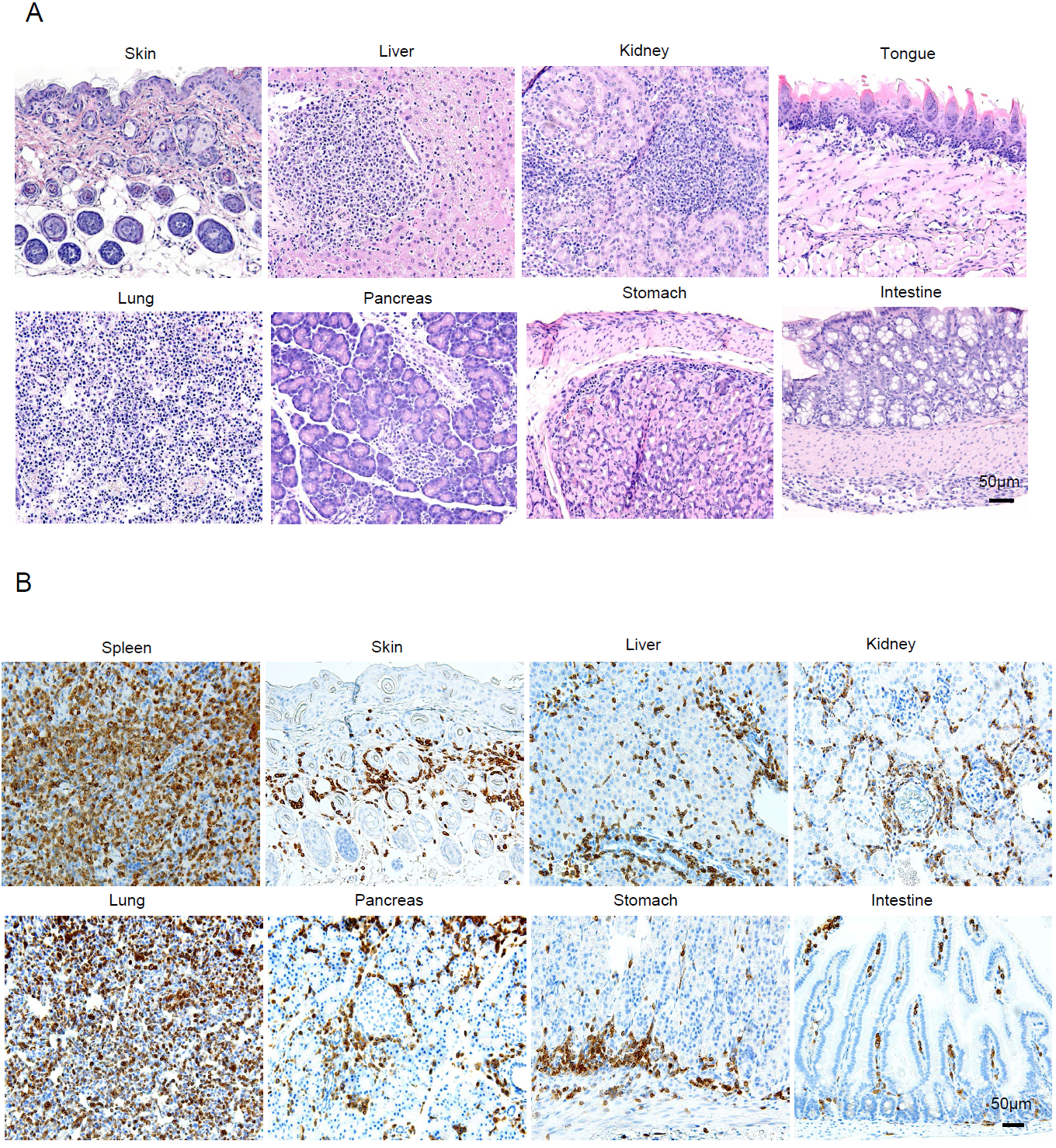
Histopathology and immunohistochemistry analyses for infiltration of human CD3^+^ cells in the organs of xenogeneic NSG recipients. A. Representative H&E in tissue sections obtained from skin, liver, kidney, tongue, lung, pancreas, stomach and intestine of GVHD mice. B. Immunohistochemical staining with anti-human CD3 shows that a high proportion of CD3^+^ cells had infiltrated various organs of human BM NSG recipients. This immunohistochemical evaluation was performed in 20 mice with similar results. Tissues were formalin fixed and paraffin embedded.

**Table 1.** Summary of pathological score in organs of xenogeneic GVHD mice

|  | Lung | Liver | intestine | skin | kidney | stomach |
| --- | --- | --- | --- | --- | --- | --- |
| no human BM (n=3) | 0 | 0 | 0 | 0 | 0 | 0 |
| BM (Donor 1) (n=3) | 3.3±0.7 | 2.7±0.3 | 2.7±0.3 | 2.7±0.3 | 2.5±0.5 | 2.5±0.5 |
| BM (Donor 2) (n=5) | 3.6±0.4 | 3.3±0.6 | 3.3±0.6 | 3.0±0 | 3.3±0.6 | 3.3±0.6 |
| BM (Donor 3) (n=7) | 3.7±0.6 | 3.7±0.6 | 3.7±0.6 | 3.7±0.6 | 3.7±0.6 | 3.7±0.6 |
| BM (Donor 4) (n=5) | 3.8±0.2 | 3.6±0.4 | 3.6±0.4 | 3.8±0.2 | 3.8±0.2 | 3.8±0.2 |

To further test the distribution pattern of the T cells, we next examined the localization of human CD4^+^ and CD8^+^ T cell subsets in the various organs of xenogeneic GVHD mice using immunofluorescence staining. As shown in Fig. 3A, CD8^+^ T cells account for over 80% of the human lymphocytes in organs such as the liver, lung, skin, kidney, and intestine. This data demonstrates that CD4^+^ and CD8^+^ T cell subsets each contribute to the phenotype, and that CD8^+^ T cells may play a dominant role. To test for infiltration of other human cell types into the organs, we performed immunofluorescence staining with anti-human CD3 and CD45 antibodies (Fig. 3B). Virtually 100% of human cells (CD45^+^) that had infiltrated the organs were T cells (CD3^+^); we found very few CD45^+^CD3^−^ cells in the tissues. Together, this data suggests that T cells are responsible for the phenotype in our GVHD model.

**Fig.3.**
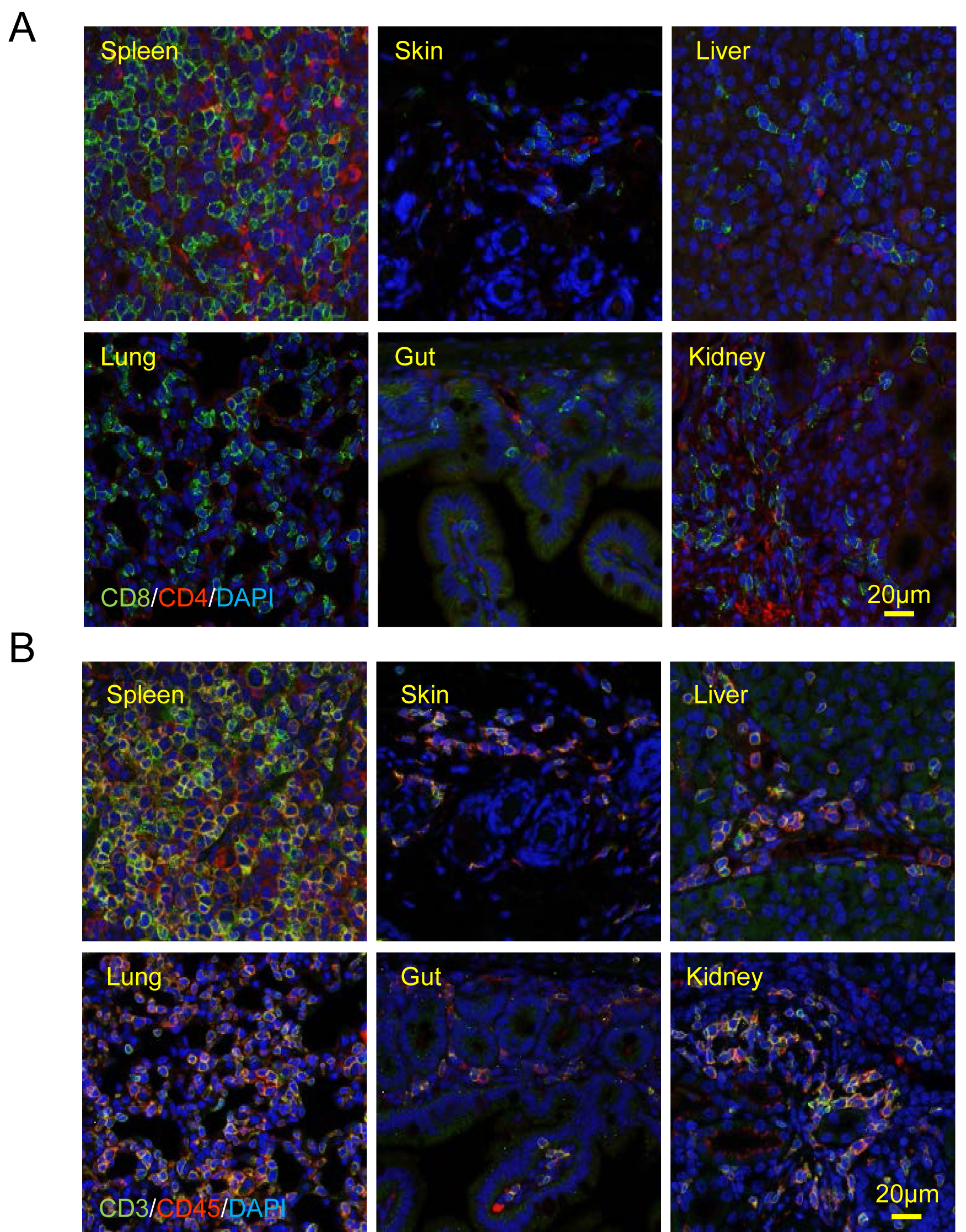
Distribution of human cells in the organs of xenogeneic NSG recipients as revealed by immunofluorescence. A. Distribution of human CD4^+^and CD8^+^cells inthe organs of xenogeneic NSG recipients. Immunofluorescence staining with anti-human CD4 and CD8 shows the distribution of CD4^+^ and CD8^+^ T cells in various organs of human BM NSG recipients. This immunofluorescence evaluation was performed in 15 mice. B. Distribution of human CD3^+^ and CD45^+^ cells in the organs of xenogeneic NSG recipients. Immunofluorescence staining with anti-human CD3 and CD45 shows the distribution of CD3^+^ and whole human CD45^+^ cells in various organs of xenogeneic NSG recipients. This immunofluorescence evaluation was performed in 15 mice. Tissues were formalin fixed and paraffin embedded. Data are representative of three independent experiments.

### Accumulation of HIF1α is critical for the maintenance of activated T cells in GVHD mice

HIF1α plays a critical role in driving T cell differentiation, metabolism and cytotoxic activity^13–15^. However, the role of HIF1α in GVHD has not been investigated to our knowledge. We first detected the expression of HIF1α by intracellular staining of cells isolated from the spleen and BM of GVHD recipients. To examine which T cell subsets were accumulating high levels of HIF1α, we stained the isolated BM and spleen cells with anti-human CD45, CD4, CD8 and HIF1α, and performed FACS analysis. We found that HIF1α was detectable at high levels in CD4^+^ and CD8^+^ T cell subsets (Fig.4A). Surprisingly, while approximately 40% of CD8 T cells in BM from the recipient mice are HIF1α^+^, more than 70% of CD8 T cells in the spleen express HIF1α (Fig. 2A). Since the spleen is normally well oxygenated, HIF1α protein must be resistant to oxygen-mediated degradation. In contrast, only 10% of CD4^−^CD8^−^ cells in the spleen express HIF1α (Fig. 2B).

**Fig.4.**
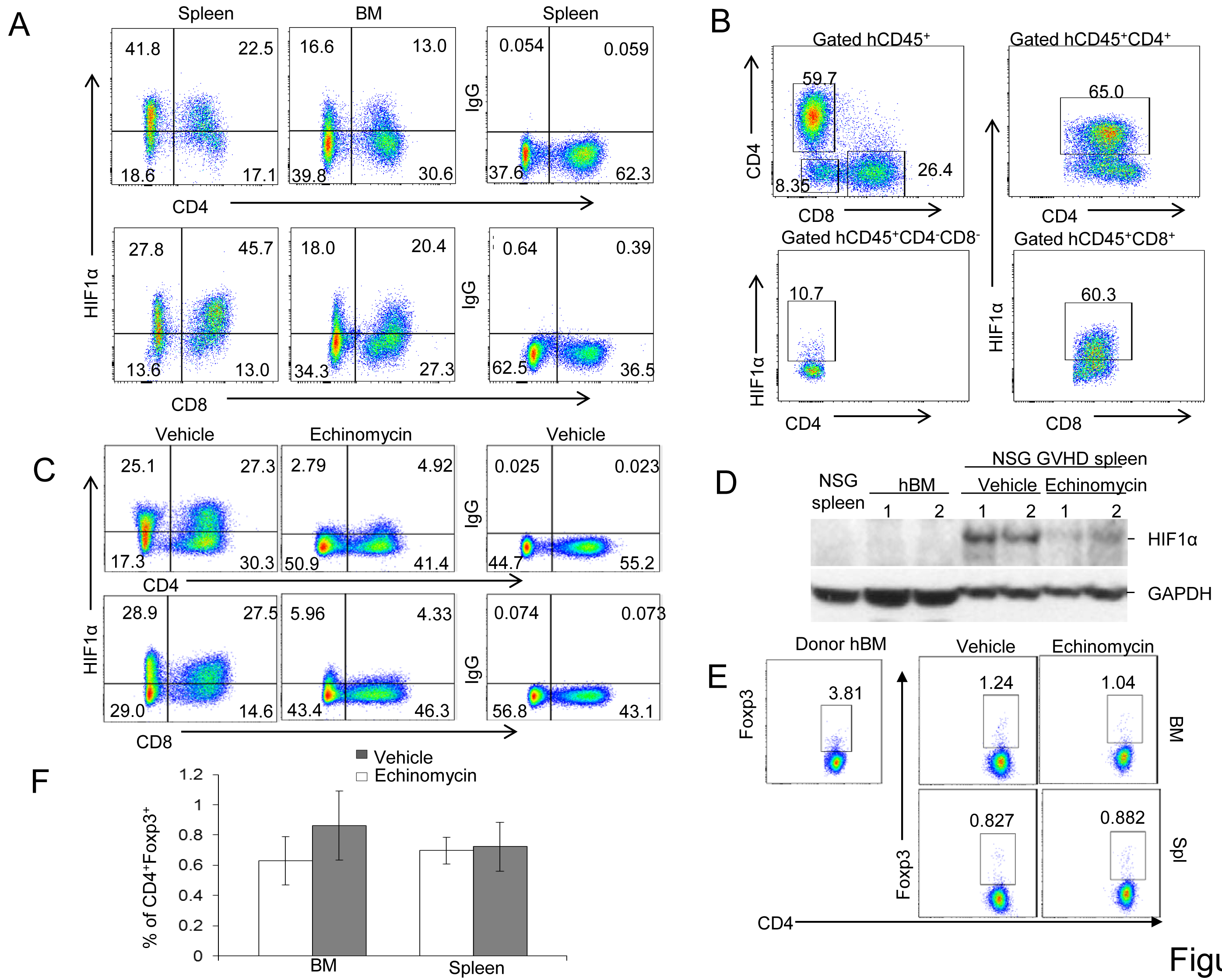
HIF1α accumulates in a high proportion of human T cells derived from spleen and BM of the recipients of human BMC: Impact of Echinomycin. A. Representative FACS plots showing the accumulation of HIF1α in T cells from the spleen and BM of GVHD mice. Spleen and BM cells were isolated from the spleens and BM of GVHD mice, stained with anti-human CD45, CD4, CD8, and then intracellular stained with anti-human HIF1α. B. Representative FACS plots showing the accumulation of HIF1α in all human cells from spleens of GVHD mice. Spleen cellswere isolated from the spleens of GVHD mice, stained with anti-human CD45, CD4, CD8, and then intracellular stained with antihuman HIF1α. FACS analysis was performed to determine the percentage of HIF1α positive cells in CD45^+^CD4^+^CD8^−^, CD45^+^CD4^−^D8^+^and CD45^+^CD4^−^D8^−^subpopulation. C. Representative FACS plots showing the percentage of HIF1α positive T cells in spleen of GVHD mice after Echinomycin treatment. Spleen cells were isolated from bone marrow chimera mice that received 5 daily doses of 10 μg/kg and one dose of 100 µg/kg of Echinomycin or vehicle before sacrifice, and stained with anti-human CD45, CD4, CD8, and then intracellularly stained with anti-human HIF1α. Data are representative of three independent experiments. D. Western blot analysis of HIF1α in spleen cells from GVHD recipients after Echinomycin treatment. Spleen cells from GVHD mice were isolated and protein lysates were subjected to immunoblot. Protein levels of HIF1α in human BM (hBM) and spleen cells from two vehicle and Echinomycin-treated GVHD mice were examined by Western blot. GAPDH served as a loading control. Data is one representative file from 3 mice and representative of three independent experiments. E and F. Echinomycin treatment does not expand human Treg in the NSG recipients of human. Spleen and BM cells were isolated from spleen and hind femur of GVHD mice treated with 10 doses of 10 μg/kg and one dose of 100 µg/kg or vehicle before sacrifice, stained with anti-human CD45, CD4, CD8, and then intracellular stained with anti-human Foxp3. Data are representative of three independent experiments. Representative FACS profiles are shown in E, while summary data (Means and SEM) are shown in F.

To test the significance of HIF1α accumulation, we treated GVHD recipient mice with Echinomycin, an inhibitor of HIF1α. Surprisingly, the HIF1α^+^ cells lost HIF1α protein overnight following a single dose of Echinomycin treatment (Fig. 4C). Since the T cell frequencies were largely unaffected, Echinomycin must have reduced expression of HIF1α rather than eliminated the HIF1α-expressing cells. Further, we isolated the spleen cells from GVHD mice and measured human HIF1α protein levels from Echinomycin-treated mice and vehicle controls by western blot analysis. HIF1α protein levels were significantly decreased in the spleen cells of Echinomycin-treated mice as compared to those of vehicle controls. Since HIF1α was absent in the human BM cells used for the GVHD induction and in NSG recipient that received no BMT (Fig. 4D), accumulation of HIF1α protein must have occurred following GVHD induction.

It has been reported that HIF1α antagonizes Foxp3 by regulating and interacting with Foxp3 to regulate T cell activation ^17,18^. To assess whether our xeno-GVHD syndrome was influenced by Treg cells via a Foxp3- HIF1α mediated regulatory circuit, we tested the effect of HIF1α inhibition on the percentage of Foxp3^+^ cells within the CD4 T cell subset derived from the spleen and BM of either Echinomycin treated or vehicle control mice. There was no apparent difference in proportion of Tregs between Echinomycin treated or vehicle control mice (Fig. 4E, F). These results suggest that HIF1α does not contribute much to maintenance of FOXP3^+^ human T cells in the NSG mice. However, since the frequency of the FOXP3^+^ T cells are much lower in the NSG mice than in the BM donor cells, it is possible that the NSG environment may not be suitable for expansion of human Treg cells.

### Echinomycin treatment protects mice against lethal GVHD

To better understand the importance of HIF1α in the therapy of GVHD, we examined the effect of inhibition of HIF1α. Newborn pups were transplanted with 0.5×10^6^ human BM cells via intra-hepatic injection. Twenty-seven days after transplantation, mice were treated with 10 µg/kg of Echinomycin for a total of 20 treatments. The therapeutic regimen is depicted in Fig.5A and an example follow up of a treated mouse is shown in Fig. 5B. As shown in Fig. 5B, at day 26 after BMT, the mouse exhibited severe skin defects, including hairlessness, inflammation and thickened and dry skin. After two weeks of treatments, the mouse regained smooth skin with visible regrowth of hair. The mouse also showed normal growth over time, although the recovery of hair was incomplete. Corresponding to improvement in clinical symptoms, the number of human leukocytes in the peripheral blood was reduced by more than 10-fold over 5 weeks (Fig. 5C). IHC analyses revealed elimination of T cells from all major organs, including skin, intestine, liver and lung (Fig. 5D).

**Fig.5.**
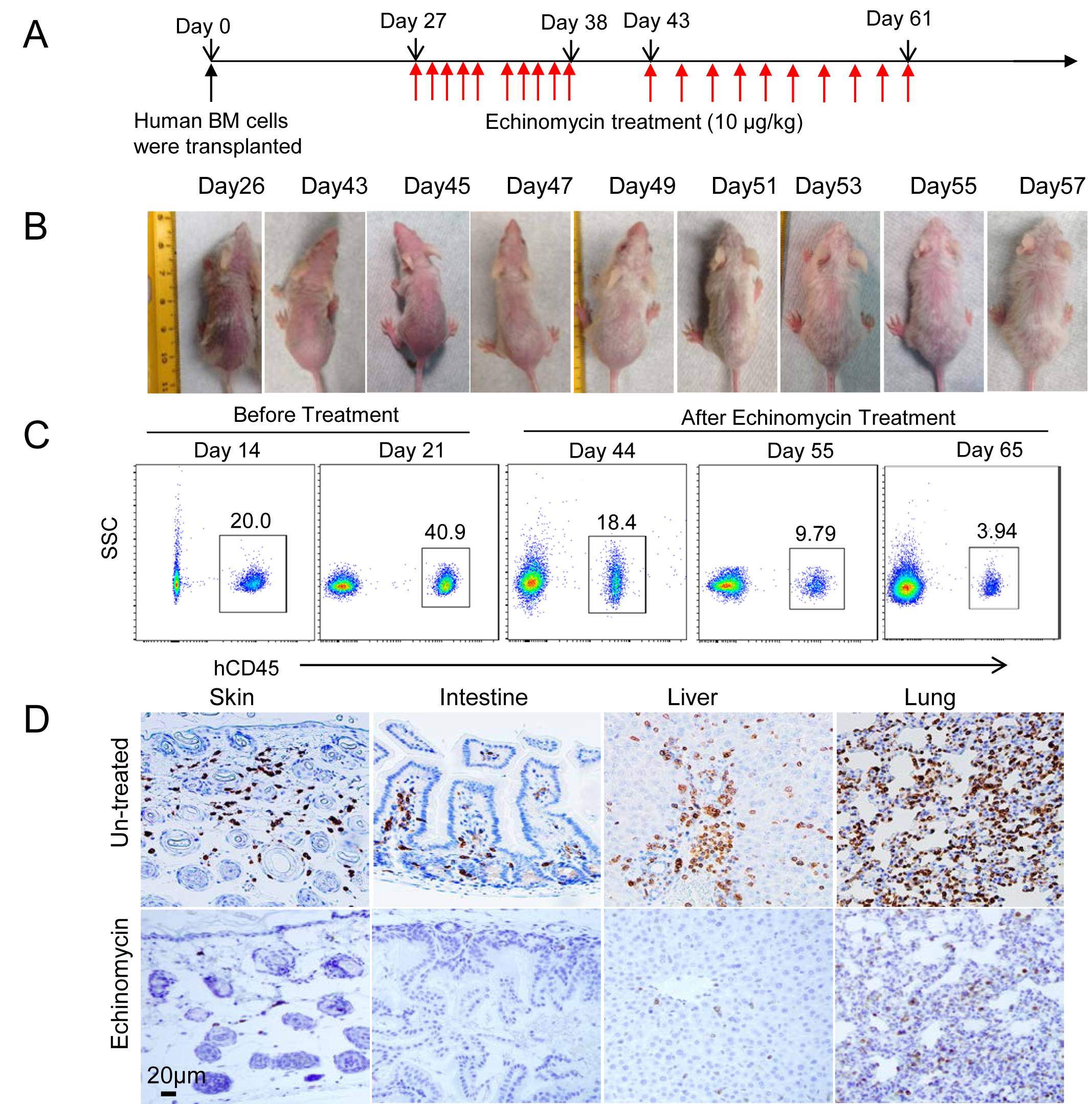
Echinomycin protects mice against GVHD and eliminates expanded human leukocytes and inflammation. A. Dosing regimen of Echinomycin treatment for humanized GVHD mice. The day that mice were transplanted with human BM cells is defined as day 0. The red arrow represents one Echinomycin treatment. The dose for each treatment is 10µg/kg via intraperitoneal injection. B. Longitudinal analysis of clinical representation of an Echinomycin-treated mice. C. Representative FACS plots showing the percentage of human CD45^+^ cells in PBL of GVHD mice before and after Echinomycin treatment. Data is one representative file from 3 mice and representative of three independent experiments. D. Immunohistochemical staining with anti-human CD3 shows a significant reduction of CD3^+^ cells in the same mouse in Fig. 5B after Echinomycin treatment at day 75 (bottom) and an untreated littermate, which died at day 55 after transplantation of human BM (Top). Data are representative of three independent experiments.

Remarkably, while all of the vehicle-treated mice died during the treatment period, none of the Echinomycin-treated mice succumbed during treatment, although the mice gradually succumb to GVHD after cessation of the drug (Fig. 6A). The median survival in the vehicle treated group is 51 days, while that of the Echinomycin treated group is 99 days (Fig. 6A). Therefore, Echinomycin extended the mouse life span even after cessation of treatment. In another experiment involving a different donor, we transplanted 0.3X10^6^ BM cells into newborn recipients and treated the recipients at day 17 after transplantation. Again, significant protection was observed as the median survival in the vehicle treated group is 50 days, while that of the Echinomycin treated group is 119 days (Fig.6B).

**Fig. 6.**
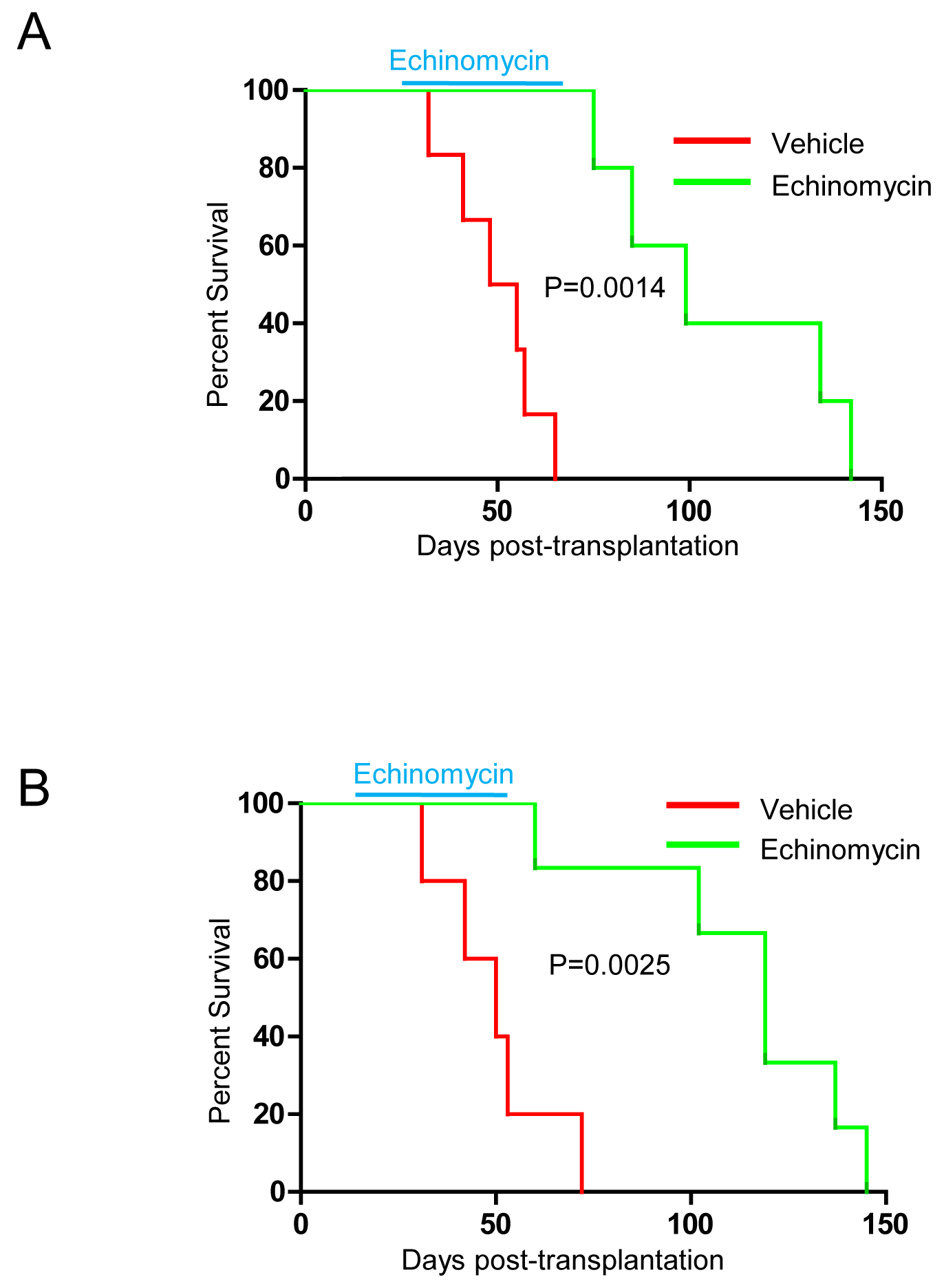
Echinomycin protects mice against lethal GVHD. A. Newborn NSG pups were transplanted with human 5X10^5^ BM cells. The mice were treated with the dosing regimen of Echinomycin, starting at day 27 as diagrammed in Figure 5A. B. Newborn NSG pups were transplanted with human 3X10^5^ BM cells. The mice were treated with the dosing regimen of Echinomycin listed in Figure 5A except of starting the treatment at day 17. Kaplan-Meier survival curve shows that recipients treated with Echinomycin displayed significantly prolonged life span compared with recipients treated with vehicle. Data are representative of three independent experiments.

## Discussion

In this study, we have developed a new humanized mouse model for GVHD by transplanting a low number of human BM cells into newborn NSG pups. We showed that the mice developed an acute GVHD syndrome with 100% mortality within 2 months. Compared with the currently used xenograft GVHD models caused by human PBL ^7-11^, our model offers several advantages. First, while the PBL-induced GVHD causes damage by inflammation primarily in the lung ^7,8^, our model recapitulates human pathology as severe inflammation was found in all clinically relevant organs, including the skin, gut, and liver. Second, the GVHD in our model is caused by T cells from human BM, which is the primary source of donor cells of HSCT in clinical practice. Third, the use of mouse pups rather than adults reduces the cost of animal care as most of the study can be completed prior to weaning.

Recent studies have shown that HIF1α plays a critical role in driving T cell differentiation, metabolism and cytotoxic activity^13-15^. T cell activation both induces and stabilizes HIF1α, leading to increased cytolytic activity of CD8^+^ T cells ^14-16^. Here we showed that HIF1α is greatly elevated during GVHD as the human T cells express high levels of HIF1α, which can be eliminated by short-term treatment with Echinomycin. Importantly, continuous treatment of Echinomycin confers protection against lethal GVHD in nearly 100% of mice. The therapeutic effect is further confirmed in that essentially all mice died after cessation of drug. While the biological impact of HIF has been well documented in models of autoimmune diseases, cancer biology, cancer immunity and viral immunity ^21^, our work is the first that extends the role for HIF into the pathogenesis of GVHD.

HIF1α is known to bind Foxp3 and repress the development of Foxp3 in vitro and in the model of experimental autoimmune encephalomyelitis (EAE) ^17^. Surprisingly, in our GVHD model, inhibition of HIF1α by Echinomycin failed to expand Tregs. We suspect that the differences may be explained by lack of necessary survival factors for human Tregs in the xenograft model. However, a recent study has suggested that over-expression of HIF1α in *Vhl*^-/-^ Tregs does not substantially alter the frequency of Tregs. Nevertheless, since HIF1α over-expression impedes the Treg function, it remains possible the surviving Tregs in our model are more active, although we consider it unlikely as the frequency is less than 1%. Regardless of whether Echinomycin has enhanced Treg function, it will likely also directly affect effector T cell function as most of them expressed HIF1α, and chronic treatment leads to depletion of most human T cells over time.

GVHD is a major barrier to HSCT in leukemia patients. Our data presented herein reveals HIF1α as a therapeutic target and Echinomycin as a potential therapeutic for the disease. Echinomycin (NSC526417) ^22^ is a member of the quinoxaline family originally isolated from Streptomyces *echinatus* in 1957 ^23^. It was brought into clinical trials by the National Cancer Institute, and multiple phase I ^24–28^ and phase II ^29–36^ trials for solid tumors have been conducted over the years. While the drug was unsuccessful for treatment of late stage solid tumors, these studies established a safety dose that is 30-50-fold higher than what is used for the treatment of experimental GVHD. Therefore, it is of great interest to test if a safe dose of Echinomycin can be useful for treating GVHD in humans.

Finally, we reported previously that Echinomycin was effective in treating experimental leukemia and lymphoma by targeting HIF1α and eliminating leukemia stem cells without adverse reactions towards hematopoietic stem cells ^20,37^. Since the majority of GVHD is observed when leukemia patients receive HSCT, Echinomycin and other HIF inhibitors may be uniquely suited for treating GVHD while reducing relapse of leukemia.

## Acknowledgments

This study was supported by the grants from the National Institutes of Health National Cancer Institute (CA171972, CA183030 (Y Liu), and CA164469 (Y Wang) and the William Lawrence & Blanche Hughes Foundation (Y Wang).

## Author contributions

Yang Liu, Pan Zheng and Yin Wang designed research, Yin Wang, Yan Liu, Christopher Baily, Chunshu Wong performed the research. Yin Wang and Yang Liu wrote the paper with input from co-authors. The authors have no conflict of interest.

